# An abundance of brain-expressed genes show ectopic activation in lung adenocarcinoma

**DOI:** 10.1101/2023.09.12.557389

**Authors:** Anna Diacofotaki, Axelle Loriot, Charles De Smet

**Affiliations:** Group of Genetics and Epigenetics, de Duve Institute, Université Catholique de Louvain; Brussels, Belgium; Group of Computational Biology and Bioinformatics, de Duve Institute, Université Catholique de Louvain; Brussels Belgium

## Abstract

Tumoral transformation processes sometimes include activation of unscheduled gene expression programs in the cancer cells. This is best exemplified by the so-called cancer-testis (CT) genes, a group of genes expressed in testicular germ cells that become activated in tumors of various somatic origins, through a process of DNA demethylation. Here, we explored the possibility that other tissue-specific gene clusters may become ectopically activated in tumors. Lung adenocarcinoma (LUAD) was used as a model, as all necessary transcriptomic datasets were available, including that of **AT2** cells, the cell-of-origin of LUAD. We found that besides CT genes, an abundant group of genes expressed in the brain (CB genes, n=63) or in both brain and testis (CBT genes, n=28) become aberrantly activated in LUAD cell lines and tissues. Interestingly, activation of CB and CBT gene clusters was also detected in various other tumor types. Most CB/CBT genes appeared to exert neuronal functions. Moreover, a significant number of them encode antigens involved in neurological paraneoplastic syndromes. Neither neuroendocrine transdifferentiation, which occurs in 10-20% LUAD, nor DNA demethylation appeared to be responsible of the ectopic activation of CB and CBT gene clusters. Instead, prediction tools and depletion experiments identified the REST repressor as a regulator of a number of CB/CBT genes. *Conclusion:* Together, our data indicate that tumor development is associated with aberrant activation of a brain gene expression program, supporting the assumption that acquisition of neuronal functions might contribute to malignancy.

## Introduction

Tumor development is driven by genetic and epigenetic alterations, which modify cancer cell functionalities by inducing changes in gene expression patterns. Such changes include the unscheduled activation of tissue-specific gene expression programs that are normally maintained silent in the cell from which the tumor originates. A well-known illustration of this is the activation of a group of testicular germline-specific genes in various tumors of somatic origin (1). Aberrant activation of these so-called **C**ancer-***T***estis (**CT**) or Cancer-Germline (**CG**) genes was found to result from the process of **DNA** demethylation that commonly affects the genome of cancer cells (2). Further analyses of transcriptomic data revealed that, besides **CT** genes, genes with specific expression in other tissues can also undergo illegitimate activation in tumors of unrelated histological types (3-7). Importantly, aberrant activation of highly tissue-restricted genes in tumors is of clinical interest, as it may offer therapeutic and diagnostic opportunities (6-8).

In most studies, cancer-activated tissue-specific genes were identified by comparing transcriptomic data of the tumor samples with that of the corresponding normal tissues. This can lead to misinterpretation, since tumors originate from a specific cell subtype, which is sometimes rare within the tissue, and whose particular gene expression profile cannot be revealed in transcriptomic data obtained from the tissue as a whole (9). There is therefore a risk that genes supposedly activated in tumors, were in fact already expressed in the cell of origin. Thus, a prior condition for identifying genes that are effectively activated during the process of tumor development is the knowledge of the cell from which the tumor originates, and the availability of expression data from this specific cell type.

In light of the limitations discussed above, we decided to re-evaluate the ectopic activation of tissue-specific gene clusters in a tumor model where the cell of origin is known. We choose lung adenocarcinoma (**LUAD**) for this exploration, as it was demonstrated that this tumor originates from alveolar type II (**AT2**) cells (10, 11), which represent about 10% of the cells that compose the lung tissue (12). Importantly, transcriptomic data of human **AT2** cells are available (13, 14). We thus performed an integrative analysis of **RNA**-seq data from **LUAD** tissues and cell lines, normal lung, and **AT2** cells, as well as of single-cell **RNA**-seq of normal tissues. Remarkably, we observed activation of a brain-specific transcriptional network in **LUAD** cells. The mechanisms underlying ectopic activation of this particular gene cluster were investigated.

## Materials and methods

### Alveolar type II (AT2) cells datasets

Fastq files of bulk **RNA**-seq experiments of **AT2** cells originating from small (n=3) and large (n=3) airways of normal lung were obtained from (13). **Q**uality control of reads was assessed using Fast**QC** v0.11.8 (Brabraham institute) and reads of low quality were discarded using Trimmomatics v0.3 (15). Reads were aligned onto hg38 genome using **HISAT2** v2.1.0, (16). FeatureCounts from the subread package v2.0.0 (17) was used for gene quantification. Expression data for each gene were averaged for small and large airway tissue samples (n=6 in total) and normalized in Transcripts Per Million (**TPM**). For **C**p**G** methylation analyses, fastq files of whole genome bisulfite-seq of **AT2** cells were obtained from the study of Zuber and colleagues (14). To assess read quality control and discard low-quality reads, Trim Galore! v0.5.0 was used (https://github.com/FelixKrueger/TrimGalore). Read alignment onto the hg38 genome and methylation calling were performed using Bismark v0.20.0 (18). Methylation percentage of **CpG** sites that were sequenced in both forward and reverse strands were averaged.

### Normal tissue samples datasets

Normalized bulk **RNA**-seq data of normal tissues were downloaded from the **GTEx** portal v7 (19). Expression data for each tissue are measured in median **TPM** of all samples per tissue analyzed. For visualization of mapped reads onto the genome of normal cerebral cortex and testis tissue samples, corresponding fastq files were downloaded from **SRA** and processed into bam files as previously described (20). **DNA** methylation datasets of normal human tissues were obtained and processed as previously described (20).

### LUAD cell lines datasets

Fastq files of bulk **RNA**-seq experiments of 26 **LUAD** cell lines were obtained from the **DBTSS** portal (21) and processed as previously described (22).

### TCGA expression datasets

Normalized bulk **RNA**-seq **FPKM** (fragments per kilobase per million) data of the **LUAD** project (23) were downloaded using the **TCGA**biolinks **R** package v2.14.1 (24). **D**ata were converted to **TPM** by dividing each **FPKM** gene expression in a sample by the sum of all gene expressions for that sample (×10^6^). **LUAD** tissues were defined as positive for the expression of a specific gene when **TPM** ≥ 2, and negative when **TPM** < 1.

### Identification of genes ectopically activated in LUAD cells (EctopEx genes)

Genes that displayed an average expression **TPM** < 0.5 in **AT2** cells and normal lung tissue samples of **TCGA** (n=58) and **GTE**x (n=578), were considered transcriptionally inactive in these cells and tissues. Among these genes, we retained those that showed an expression **TPM** ≥ 2 in at least 5% of the total number of **LUAD** tissue samples available from **TCGA** (n=510). Finally, we only retained the genes that were activated in at least one **LUAD** cell line and repressed in at least two **LUAD** cell lines of the 26 **LUAD** cell lines available from **DBTSS**. Activation in these cells was defined as **TPM** ≥ 2, and repression as **TPM** ≤ 0.1.

### Characterization of tissue-specific gene clusters among EctopEx genes

The Ward.**D** method and Euclidean distance were used for unsupervised hierarchical clustering analyses of EctopEx gene expression in **GTE**x samples. **C**ancer-**B**rain (**CB**) genes were defined as follows: they showed either an expression **TPM** ≥ 2 in at least one brain tissue from those available in **GTE**x and **TPM** < 1 in all other tissue samples, or they showed a median expression ≥ 1 **TPM** in at least one brain tissue and in maximum one other tissue (other than the testis). Genes that displayed a median expression **TPM** ≥ 1 in at least one of the brain tissues available from **GTE**x and in testis, and exhibited a median expression **TPM** < 1 in all other tissues, were defined as Cancer-Brain/Testis (**CBT**) genes. Genes that showed a median expression **TPM** ≥ 2 only in testis of **GTE**x, or a median expression **TPM** ≥ 1 in testis and in maximum one other tissue (except brain), were defined as **C**ancer-**T**estis (**CT**) genes. For establishing the expected number of **CB, CBT**and **CT** genes, we first applied the selection procedure described here above to all the genes that were found to be initially silent in normal lung and **AT2** cells, thereby identifying total numbers of brain-, brain/testis-, and testis-specific genes. These numbers were then multiplied by the overall proportion of silent genes that we found to be activated in **LUAD**. For evaluation of potential enrichment of molecular functions and biological process subcategories of Gene Ontology annotations, the clusterProfiler R package v4.0.2 was used (25). To accurately identify the TSS of each **CB** and **CBT**genes and confirm their activation in **LUAD** cells, BAM files of cerebral cortex, testis tissue samples and the 26 **LUAD** cell lines, were visualized in the **IGV** software (26).

### Single-cell RNA-seq analyses

Normalized single cell **RNA**-seq expression data were downloaded from the Human Protein Atlas (HPA) v20.1 (27). Expression data from different clusters of the same cell type were averaged to obtain a mean expression of each gene per cell type. Data are expressed in p**TPM** (transcripts per million of protein coding genes).

### Immunohistochemical data

IHC images of normal brain, normal lung and lung adenocarcinoma tissues were obtained from the Human Protein Atlas (HPA) v20.1. The most reliable IHC data were selected on the basis of the following criteria: positive staining was confirmed by HPA, and correlated with **RNA**-seq gene expression profiles of **GTE**x and HPA consortia. HPA codes for antibodies and ID numbers of tissue sections illustrated in this study were the following: HPA023099 and ID3618 (Cerebellum), ID4797 (Lung), ID4923 (**LUAD**) for CALB1; HPA049148 and ID3731 (Cerebral cortex), ID2208 (Lung), ID4692 (**LUAD**) for DPF1; HPA067549 and ID3740 (Cerebral cortex), ID1678 (Lung), ID4208 (**LUAD**) for OPRK1.

### i-cisTarget analysis for transcription factor binding

The i-*cis*Target webtool was used with default parameters and ENSEMBL gene names as input. We selected the following ChIP studies against **REST**: H**CT**116 (ENCFF001UEC), MCF7 (ENCFF001UNI) and PANC1 (ENCFF001UOD). When multiple conserved regions of the same gene displayed an enrichment for **REST**, enrichment ranking of the region closest to the TSS was considered.

### RNA-seq of C2 dermal fibroblasts transfected with sh-REST

FastQ files of C2 cells transfected with a control or anti- **REST** sh-**RNA** were obtained from the study of Drouin-Ouellet (28). FastQ files were processed as described above for **AT2** cells. DESeq2 R package v1.32.0 was used for differential expression analysis, with counts generated by Featurecounts as input.

### Comparison of gene expression levels in tumor cell lines

The expression of **CB** and **CBT**genes was compared with that of ***REST*** in tumor cell lines listed in the Cancer Cell Line Encyclopedia (**CCLE**), through the use of the Depmap portal (29). Tumor cell lineages classified as “central nervous system”, “peripheral nervous system”, and “unknown” were excluded from the analysis. **CB** and **CBT**gene expression levels (log2(**TPM**+1)) were compared in ***REST***^low^ (log2(**TPM**+1) <2, n=78) and ***REST***^high^ (log2(**TPM**+1) ≥2, n=1204) cell lines. Median level of ***REST*** expression (log2(**TPM**+1)) among all cell lines was 3.2.

### Correlation of CB/CBT expression in LUAD with overall survival

The GEPIA2 portal was used to evaluate correlation of **CB** and **CBT** gene expression with overall survival of **LUAD** patients (30). For all genes, cutoffs were set at 90% for the high group and 50% for the low group. Kaplan-Meier curves were downloaded when Logrank p-value was<0.01.

### Statistical analyses and graphical representations

Statistical analyses were performed in R 4.1.0 (http://www.R-project.org). p-values were adjusted using the Benjamini-Hochberg method and a p-adjusted value < 0.05 was considered statistically significant. Graphs were generated with the following R packages: ggplot2 v.3.3.5, ComplexHeatmap v2.8.0 (31), and clusterProfile v4.0.2.

## Results

### Identification of tissue-specific gene clusters with ectopic expression in LUAD

In order to uncover tissue-specific gene clusters showing ectopic activation in **LUAD** cells, we integrated **RNA**-seq datasets of **AT2** cells, normal tissues (**GTE**x), **LUAD** cell lines (**DBTSS**) and **LUAD** tissue samples (**TCGA**), and applied the following selection workflow (**Fig. 1A**). We first selected genes that are silent (**TPM** < 0.5) in **AT2** cells and in normal lung. Among these genes, we retained those that were activated (**TPM** ≥ 2.0) in at least 5% of the available **LUAD** tissue samples. Finally, in order to isolate genes with ectopic activation in cancer cells, rather than in nonmalignant cells of the tumor microenvironment, we selected the genes that were also activated in at least one out of the 26 analyzed **LUAD** cell lines. Our procedure resulted in the selection of 1051 genes that were silent in **AT2** cells and normal lung, and showed ectopic activation in **LUAD** tissues and cell lines (“EctopEx” genes, **Fig. 1A**).

**Figure 1:**
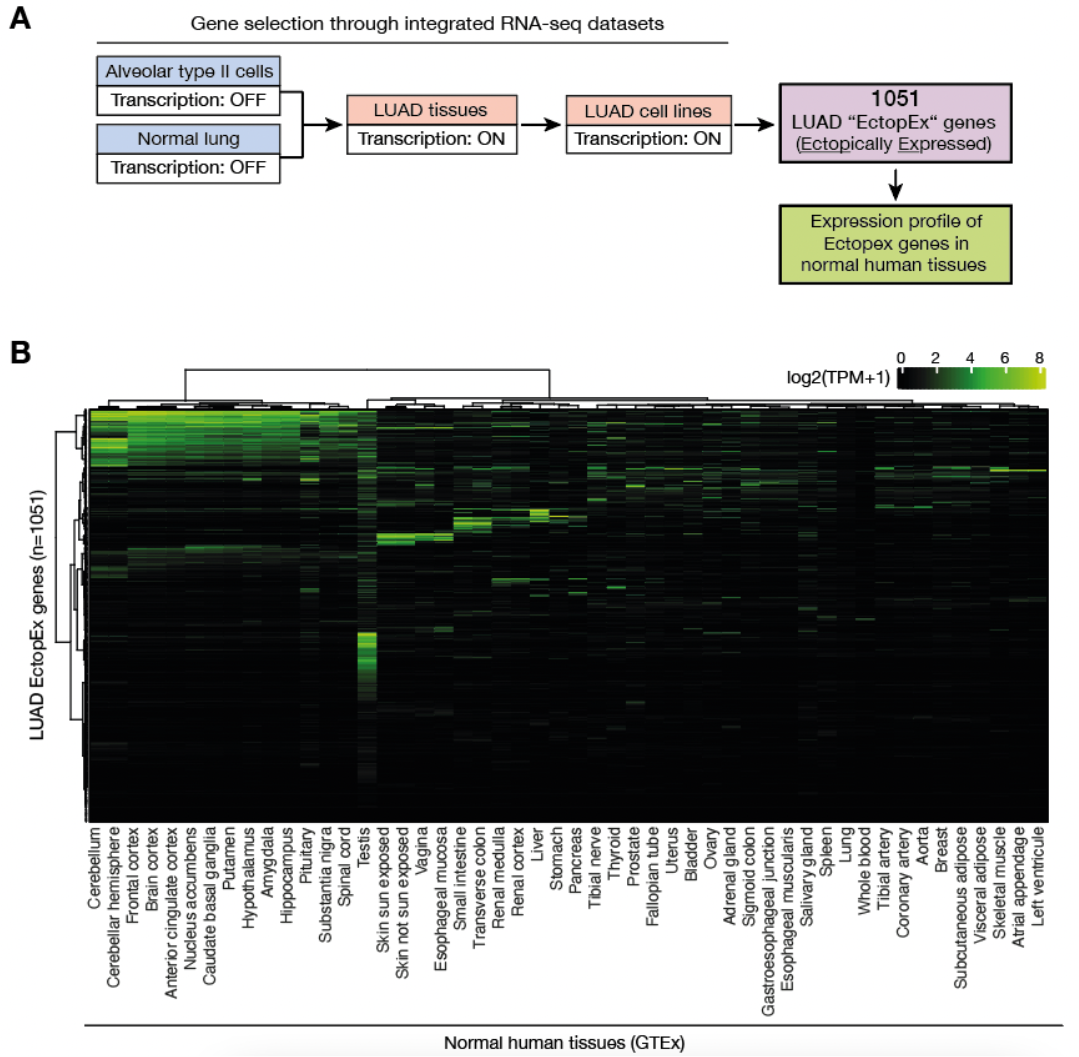
LUAD cells show aberrant activation of genes belonging to tissue-specific expression clusters. **A)** Workflow of the analysis of **RNA**-seq data for the selection of genes ectopically activated in **LUAD** cells (EctopEx genes). **B)** EctopEx gene expression in a panel of normal tissues available from **GTE**x was clustered using Ward’s method and Euclidean distance. Expression data are depicted in log2 transformed median **TPM** + 1.

In the next step, we searched to determine the expression profile of **LUAD** EctopEx genes among healthy tissues by examining **RNA**-seq datasets from a panel of normal tissue samples (**GTE**x; **Fig. 1B**). Unsupervised hierarchical clustering of gene expression in these tissues was applied, and revealed several groups of EctopEx genes showing congruent expression in defined tissues. First, there was a group of genes that showed highest expression in the testis. Closer examination of the genes in this category revealed the presence of multiple genes of the cancer-testis (**CT**) group of genes. Another group of EctopEx genes showed maximum expression in the esophagus, skin and vagina, and was therefore reminiscent of the stratified epithelium (SE) gene cluster that we described previously (20). A third group of genes with predominant expression in colon and small intestine, and occasionally in liver, stomach and/or pancreas, corresponded to the previously identified gastrointestinal (GI) gene cluster (7, 20).

Intriguingly, a large cluster of EctopEx genes displayed highest expression in brain tissues, including the hypothalamus, pituitary gland, cerebellum, and/or basal ganglia (putamen, nucleus accumbens or caudate nucleus) (**Fig. 1B**). Several of these genes appeared to be simultaneously expressed in testis. The remaining EctopEx genes displayed disparate patterns of expression among the analyzed tissues, and could therefore not be classified into a defined cluster, or showed expression levels that were below the fixed threshold (**TPM** < 2) in all tissues. Together, these results revealed that, besides the previously identified **CT**, SE and GI genes, a large cluster of genes with high expression in the brain become ectopically activated in **LUAD** cells.

### Brain and Brain/Testis genes with ectopic activation in LUAD

To get further insight into the group of brain-expressed genes showing ectopic activation in **LUAD**, we first specified a list of EctopEx genes belonging to this category by applying the following criteria: genes had to be expressed in at least one brain tissue sample (**GTE**x) and could also be expressed in maximum one other tissue. This resulted in the selection of 91 EctopEx genes with preferential expression in the brain. Strikingly, a significant proportion of these genes (28/91) showed expression in testis in addition to the brain. We therefore defined two gene subgroups: the “Cancer-Brain” (**CB**) genes showing expression in the brain but not in testis (n=63), and the “Cancer-Brain/Testis” (**CBT**) genes displaying expression in both brain and testis (n=28; **Fig. 2A**). For comparison, identification of Ectopex genes with preferential expression in testis, but not brain, resulted in the isolation of 184 “Cancer-Testis” (**CT**) genes.

**Figure 2.**
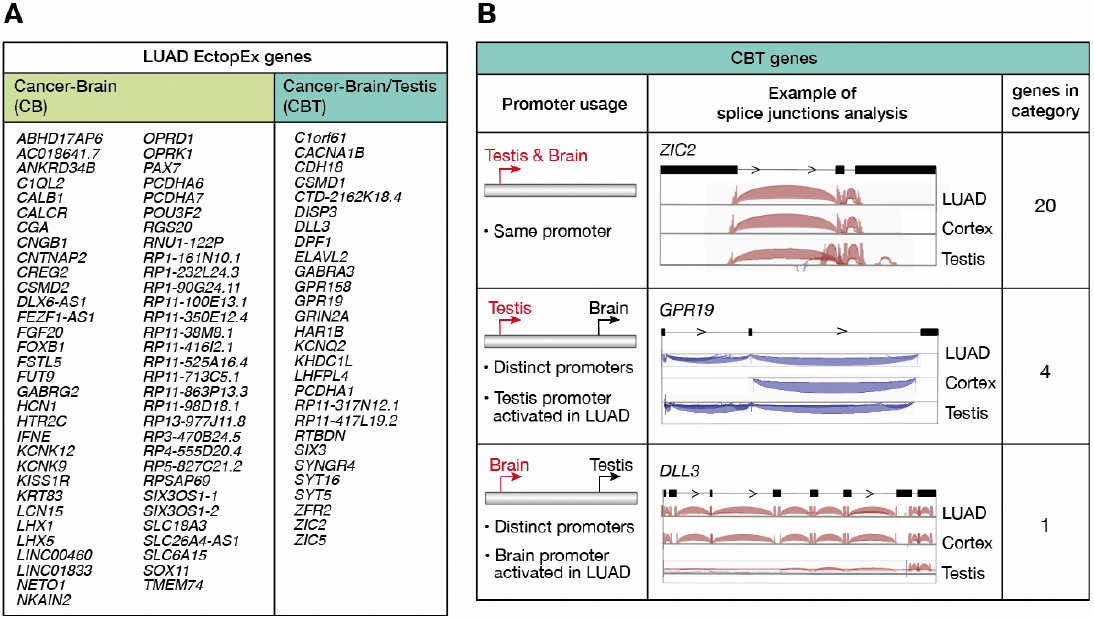
Characterization of Cancer-Brain (CB) and Cancer-Brain/Testis (CBT) genes, and identification of their promoter usage. **A)** Lists of **LUAD** Ectopex genes in the **CB** or **CBT** category were defined according to their tissue-specificity of expression (**GTE**x **RNA**-seq). **B) CBT**transcripts were analysed with the Integrative Genomics Viewer (**IGV**) to define transcriptional start sites (TSS) usage in **LUAD** cell lines, brain cortex, and testis. The table depicts the different cases of TSS usage. Left: schematic representation of possible **CBT** promoter usage (broken arrows depicting TSS). Middle: screenshot of **IGV** views of representative **CBT** transcripts (black boxes: exons; connecting lines: introns; and splice junctions of **RNA**-seq reads represented in red or blue according to the orientation of the gene on the chromosome). Right: number of **CBT** genes with corresponding promoter usage.

One possible explanation for the bimodal expression of **CBT** genes could be the usage of alternative promoters, one that is active in brain and the other in testis. This was previously described in the *GABRA3* locus, where it was found that transcriptional activation of the gene in tumor cells originated specifically from the testis-specific promoter (32). We therefore decided to precisely determine the transcription start sites (TSS) used by **CBT** genes in normal brain cortex, testis and **LUAD** cell lines by visualizing **RNA**-seq reads with the Integrative Genomics Viewer (**IGV**). For the majority of **CBT** genes that could be analyzed (20/25), we observed that transcription in the brain, testis and **LUAD** cells originated from one same TSS. For only five **CBT** genes, we observed alternative usage of different TSS in brain and testis: activation in **LUAD** cells occurring at the testis-specific TSS in four cases, and at the brain-specific TSS in one case (**Fig. 2B**).

### A majority of CB and CBT genes display neuronal expression and function

As the brain tissue is composed of a heterogeneous cell population, it was important to determine if **CB** and **CBT** genes are expressed in a particular cell type. To this end, the expression of **CB** and **CBT** genes was examined across single cell **RNA**-seq data derived from a brain tissue sample (HPA). The results showed that most **CB** and **CBT** genes (70%) are expressed in neuronal cells, and only a minority in supporting glial cells or in microglial cells (**Fig. 3A**). Consistently, examination of cellular functions associated with **CB** and **CBT** genes revealed enrichment for biological processes such as regulation of membrane potential and neural precursor cell proliferation (**Fig. 3B**). Further analysis of Panther annotations indicated that proteins encoded by **CB** and **CBT** genes exert a variety of molecular functions, including receptor signaling, ion transport, cell adhesion, and transcriptional regulation.

**Figure 3:**
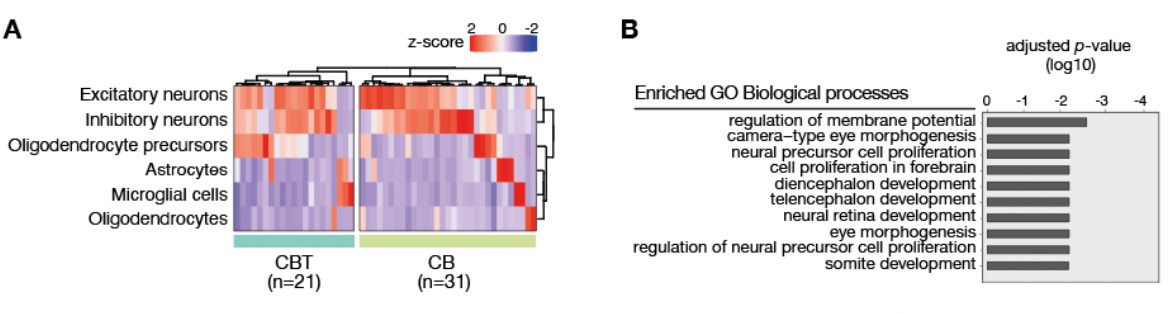
Most CB and CBT genes are expressed in neuronal cells. **A)** Single-cell **RNA**-seq data of brain tissue available from HPA were explored to determine cell type-specificity of **CB** and **CBT** genes. Gene expression was clustered using Ward’s method and Euclidean distance in the available cell types of the brain. Data are depicted using a z-score of protein coding transcripts per million. **B)** Results of enrichment analysis of **CB** and **CBT** genes using “biological process” subcategories of Gene Ontology annotation. Top 10 enrichment scores are depicted (p-adjusted <0.05).

### Frequent and robust activation of CB and CBT genes in LUAD samples

The frequency of activation and level of expression of **CB** and **CBT** genes in **LUAD** tissues were investigated in the **TCGA** datasets. The frequency of activation of individual **CB** and **CBT** genes (**TPM** ≥2) among **LUAD** tissue samples (n=510) ranged from 5% to 70%, with an average activation frequency of 13.8%. In **LUAD** tissues where **CB** or **CBT** genes were activated, the mean level of expression was 11.4 **TPM** (**Fig. 4A, B**). This was similar to what was observed for testis-specific (**CT**) genes in these tissues, with an average activation frequency and a mean expression level of 13.6% and 13.2 **TPM**, respectively. Unsupervised hierarchical clustering did not reveal preferential activation of **CB**/**CBT** genes in subgroups of **LUAD** samples, but suggested a more general mechanism of de-repression of these genes (**Fig. 4A**). Immunohistochemical data, which were available for several **CB** and **CBT** proteins (HPA), confirmed aberrant expression at the protein level in **LUAD** tissues (**Fig. 4C**). Together, these observations revealed frequent and robust transcriptional activation of **CB** and **CBT** genes in **LUAD** tissue samples.

**Figure 4.**
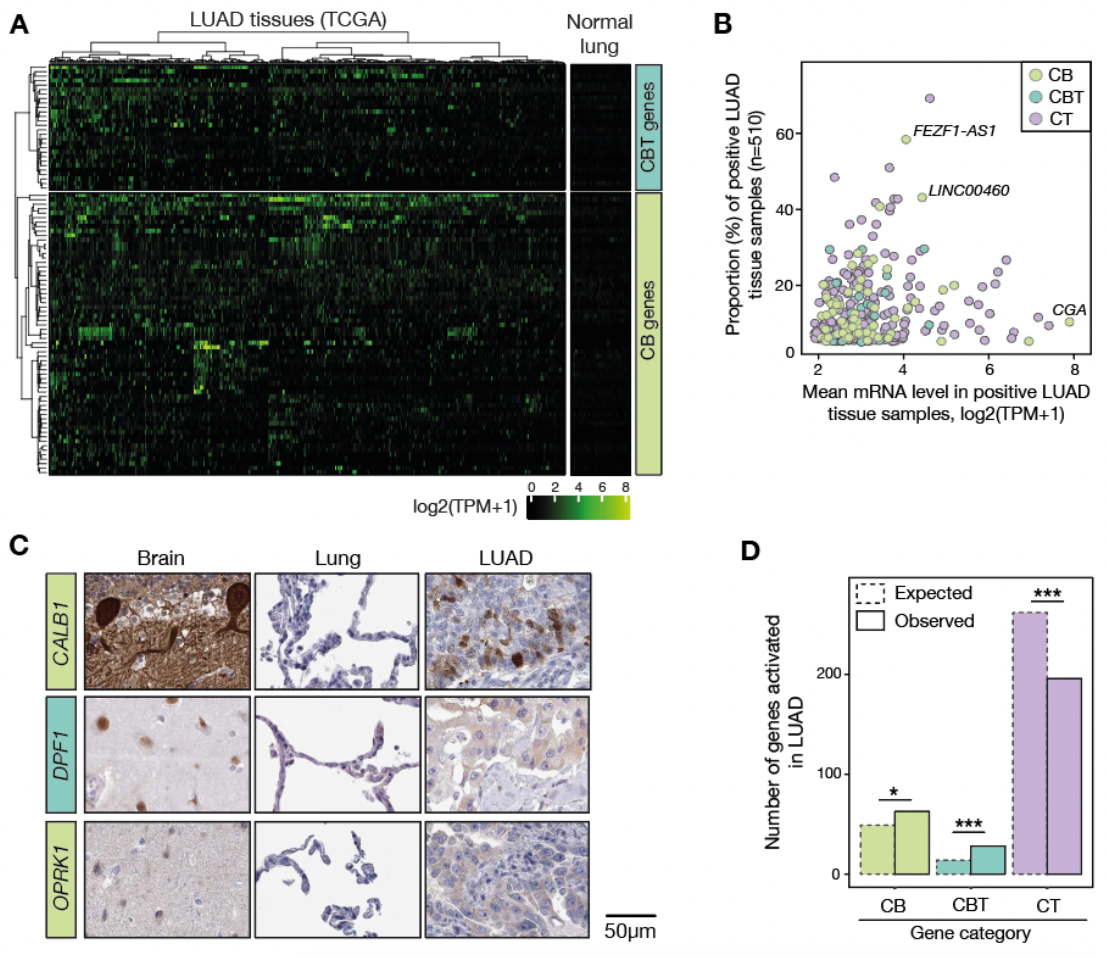
Frequent and robust activation of CB and CBT genes in LUAD tissues. **A)** Heatmap depicting m**RNA** expression levels (log2 **TPM**+1) of **CB** and **CBT** genes in **LUAD** tissues and normal matching tissues (**TCGA**). Unsupervised hierarchical clustering was applied to **LUAD** samples. **B)** Activation frequencies (**TPM** ≥2) and mean expression levels of **CB, CBT**and **CT** genes in **LUAD** samples were computed. **C)** Immunohistochemical data for **CB**/**CBT** proteins that were available at the HPA are presented. HPA scored these proteins positive in normal brain and **LUAD**, but negative in normal lung. **D)** Observed numbers of **CB, CBT**and **CT** genes with ectopic expression in **LUAD** were compared to their expected numbers, which were inferred from their proportions among the genes initially identified as not being expressed in normal lung and **AT2** cells. (Chi-squared test, ***: p-value < 0.001, *: p-value < 0.05).

The substantial amount of **CB** and **CBT** genes that we found to become activated in **LUAD** raised the possibility that there may be positive selection for activation of brain-expressed genes in the cancer cells. However, previous examination of the human transcriptome revealed that many genes display preferential expression in brain and/or testis (33). Therefore, another plausible explanation for the abundance of **CB** and **CBT** genes in **LUAD** was that it merely reflected the preponderance of human genes with these tissue-specificities. To decide between these two hypotheses, we compared the observed numbers of **CB** and **CBT** genes with their expected numbers, which were calculated on the basis of the relative proportion of brain- and brain/testis-specific genes among the genes that our selection procedure initially classified as repressed in normal lung and **AT2** cells. The results showed that the observed numbers of **CB** and **CBT** genes exceeded their expected numbers, thereby supporting selective activation of these groups of genes in **LUAD** (**Fig. 4C**). Of note, the results did not support such selectivity for the testis- specific **CT** genes, which were instead less abundant than expected (**Fig. 4C**).

### Activation of CB/CBT genes in LUAD does not coincide with neuroendocrine differentiation

Previous studies have reported that 10 to 20% of **LUAD** tumors show evidence of neuroendocrine differentiation, and that upregulation of the ASCL1 transcriptional regulator is a key factor in this phenotypic conversion (34-36). We therefore decided to determine if activation of **CB** and **CBT** genes in **LUAD** coincides with neuroendocrine differentiation. To this end, we determined the profile of *ASCL1* activation in **LUAD** tissues of **TCGA**. *ASCL1* activation was detected in 21% of **LUAD** tissues (**TPM** ≥ 2), and was correlated with the expression of the synaptophysin gene (*SYP*), a common surrogate marker of neuroendocrine differentiation (**Fig. 5A**). We then computed a Pearson correlation coefficient between the expression of *ASCL1* and that of **CB** and **CBT** genes. The results showed that the expression of only a few **CB** (5/63) and **CBT** (1/28) genes displayed significant positive correlation with *ASCL1* expression in **LUAD** tissues (ρ > 0.4, *p*-adjusted < 0.0001; **Fig. 5B**). The gene with the highest correlation coefficient was *DLL3*, a well-known target of ASCL1 (37, 38). Apart from these few genes, the vast majority of **CB** and **CBT** genes (93%) did not show significant correlation with *ASCL1* expression, thereby excluding the assumption that their activation in **LUAD** is a direct consequence of the process of neuroendocrine differentiation.

**Figure 5:**
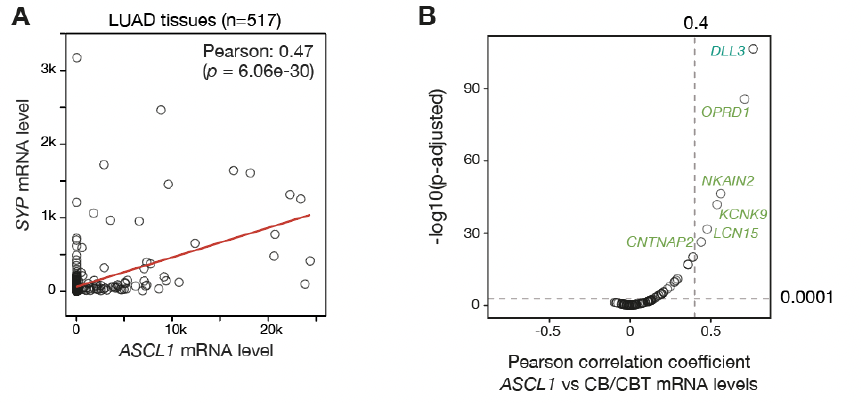
Activation of most CB/CBT genes in LUAD is not correlated with neuroendocrine differentiation. **A)** m**RNA** expression levels (**TPM**) of *ASCL1* (neuroendocrine differentiation driver) and *SYP* (neuroendocrine differentiation marker) in **LUAD** were compared, and the Pearson correlation was calculated. **B)** m**RNA** levels of each **CB** /**CBT** gene were compared with that of ASCL1, and Pearson correlation coefficients and adjusted *p*-values were calculated. Genes with correlation coefficient ≥ 0.4 and *p*-value ≤ 0.0001 are indicated.

### Up-regulated expression of CB and CBT genes in other tumor types

Having demonstrated that **CB** and **CBT** genes become activated in **LUAD** tumors, and that this is not merely associated with neuroendocrine differentiation, we next explored the possibility that these genes might also be upregulated in other tumor types. To this end, we compared the mean expression levels of **CB** and **CBT** gene sets in **TCGA** tumor collections and in corresponding normal tissues, using the GEPIA2 web server (30). Interestingly, a variety of tumor types displayed higher expression of **CB** and **CBT** gene sets when compared with corresponding normal tissue samples (**Fig. 6**), thereby suggesting that tumoral activation of such genes is not restricted to **LUAD**.

**Figure 6:**
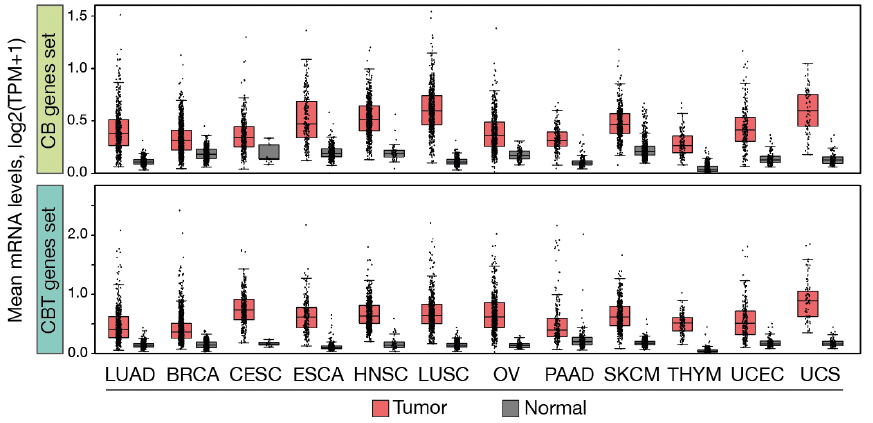
CB/CBT gene cluster are upregulated in other tumor types. Boxplot representation of the mean level of m**RNA** expression of **CB** and **CBT** gene sets in the indicated tumor types (**TCGA**) and corresponding tissue samples (**TCGA** and **GTE**x). Analysis was performed through the **GEPIA2** portal. Each dot represents the combined level of expression of all genes of the **CB** or **CBT** set in a sample. Analyzed tumor types include lung adenocarcinoma (**LUAD**), breast invasive carcinoma (**BRCA**), cervical squamous cell carcinoma and endocervical adenocarcinoma (CESC), esophageal carcinoma (ESCA), head and neck squamous cell carcinoma (**HNSC)**, lung squamous cell carcinoma (LUSC), ovarian serous cystadenocarcinoma (OV), pancreatic adenocarcinoma (**PAAD**), skin cutaneous melanoma (**SKCM**), thymoma (**THYM**), uterine corpus endometrial carcinoma (UCEC), uterine carcinosarcoma (UCS).

### CB/CBT genes are not controlled by DNA methylation

Considering the previously described role of DNA demethylation in ectopic activation of tissue-specific gene clusters in tumors (2, 20), we sought to determine if DNA methylation is involved in the regulation of **CB** and **CBT** genes. By definition, genes that become activated by DNA demethylation in tumors have their promoter highly methylated in all normal tissues where they are not expressed. We therefore examined the DNA methylation level of **CB** and **CBT** gene promoters in a panel of human tissues (ENCODE), and in **AT2** cells. For comparison, the mean promoter DNA methylation level of **CT** genes was also analyzed, as it is known that many of these genes rely on DNA hypomethylation for activation in tumors (2). The results showed that, whereas **CT** genes exhibit high promoter methylation levels in all tissues (lower in testis, which contains expressing germ cells), **CB** and **CBT** genes have a promoter that is generally unmethylated, with no difference between the expressing brain and the other non-expressing tissues (**Fig. 7**). We conclude from these observations that DNA methylation is not generally involved in controlling the tissue-specific expression of **CB** and **CBT** genes, and that activation of these gene clusters in tumors can therefore not be ascribed to a process of promoter DNA demethylation.

**Figure 7.**
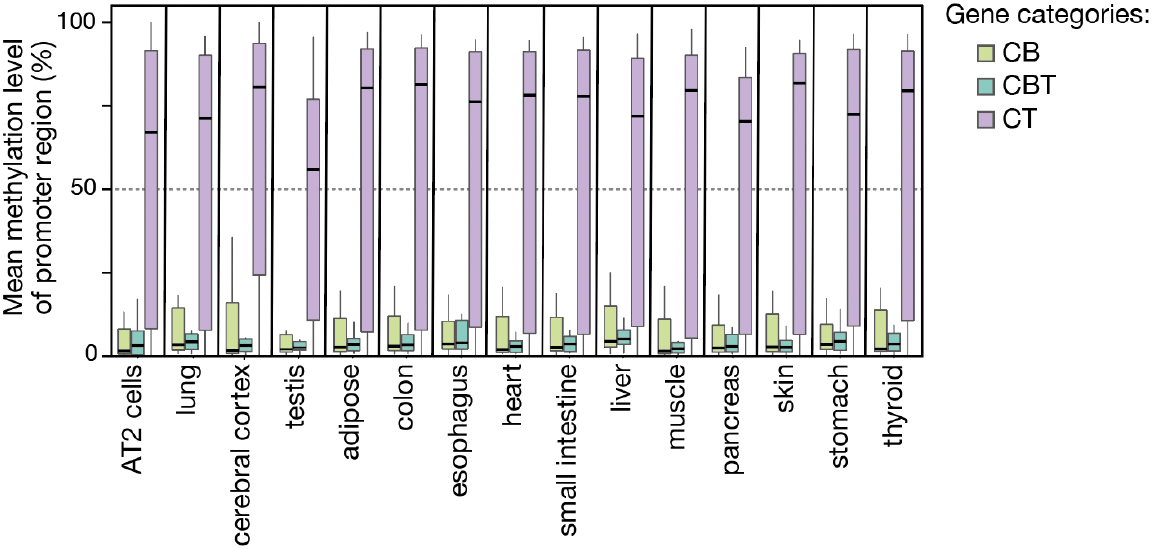
Tissue-specific expression of CB and CBT gene sets is not associated with differential promoter DNA methylation. Whole genome bisulfite sequencing data from normal human tissues and **AT2** cells were analyzed to determine the combined level of promoter DNA methylation of all genes included in either the **CB, CBT** or **CT** gene sets. Boxplot representation of the results.

### The REST repressor is involved in the regulation of a subset of CB and CBT genes

To get further insight into the mechanisms leading to the ectopic activation of **CB** and **CBT** genes in tumor cells, we decided to look for transcription factors possibly involved in their regulation. To this end, we used the i-*cis*Target online portal, which compiles ChIP-seq data and transcription factor binding site motifs in conserved genomic regions. Examination of **CB** and **CBT** genes with this prediction tool revealed highest enrichment scores for the **REST** transcriptional repressor. Indeed, a number of **CB** (n=12) and **CBT** (n=8) genes were identified as highly-ranked targets of **REST** in ChIP-seq experiments (**Fig. 8A**). The **REST** repressor is involved in the silencing of neuronal genes in non-neuronal cells (39). Consistently, all **CB** and **CBT** genes that were identified as **REST** targets were among the genes that displayed preferential expression in neuronal cells in the brain. To verify the involvement of **REST** in controlling the expression of **CB** and **CBT** genes, we analyzed **RNA**-seq data obtained from adult dermal fibroblasts producing either a control or an sh**RNA** directed against ***REST*** (28). For a subset of **CB** and **CBT** genes we observed significant m**RNA** increase following ***REST*** depletion compared to the control condition (**Fig. 8B**). As expected, this transcriptional induction was not observed for testis-specific **CT** genes.

**Figure 8:**
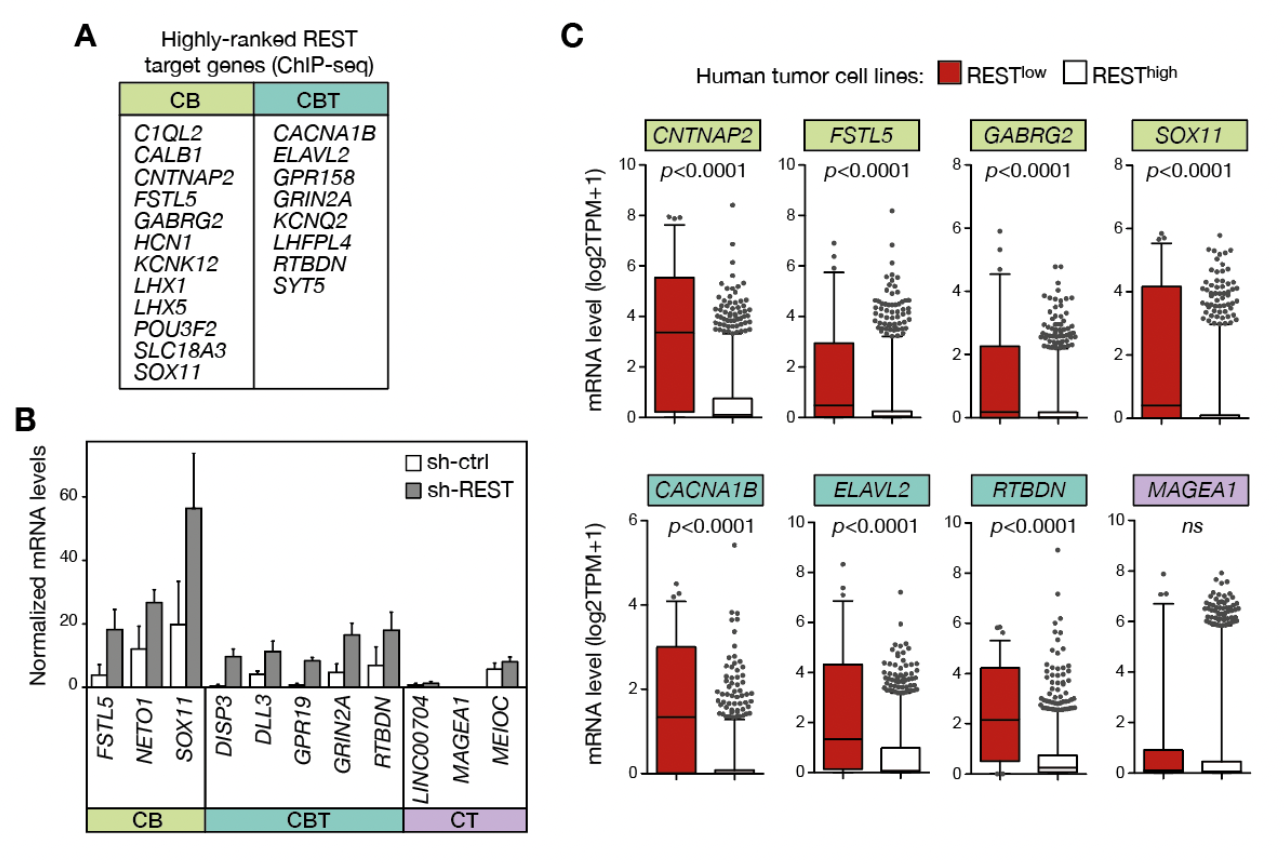
REST repressor complex is involved in the regulation of a subset of CB and CBT genes. **A)** List of **CB** and **CBT** genes showing binding of the **REST** repressor (ChIP-seq) according to the i-*cis*Target webtool. **B) RNA**-seq data (normalized counts) of adult dermal fibroblasts transfected with either control (sh-ctrl) or ***REST***-targeting sh**RNA**s (sh-**REST**) were analyzed. Expression levels of **CB**/**CBT** genes displaying significant induction upon **REST** depletion (*p*-value < 0.05) are depicted. The data of three **CT** genes (not expected to be regulated by **REST**) are shown for comparison. **C)** m**RNA** levels of **CB**/**CBT** genes were compared in ***REST***^low^ (n=78) and ***REST***^high^ (n=1204) tumor cell lines of various origins (excluding the nervous system). Boxplots (Whiskers, 5-95 percentile) of select **CB**/**CBT** genes are depicted, together with that of *MAGEA1* used as a **REST**-independent control gene. *P* values were calculated (Mann Whitney tests).

Downregulated expression of ***REST*** has been previously reported in several types of non-neuronal human tumors (40, 41). We therefore examined if ectopic activation of **CB** and **CBT** genes in tumor cells correlated with ***REST*** downregulation. To this end, **RNA**-seq data from the various tumor cell lines included in the Cancer cell line encyclopedia (except those originating from tumors of the central or peripheral nervous system) were examined. The data revealed that all **CB** and **CBT** genes that were predicted to be targeted by the **REST** repressor (listed in **Fig. 8A**), except two (*LHX1* and *GRIN2A*), displayed preferential upregulation in tumor cell lines with downregulated ***REST*** expression (**Fig. 8C**). A number of tumor cell lines where ***REST*** m**RNA** expression was not decreased nevertheless showed upregulated expression of **CB** and **CBT** genes (outliers in **Fig. 8C**), suggesting the existence of alternative mechanisms of induction of these genes in tumor cells.

### Clinical parameters associated with CB/CBTgene expression in tumors

In order to evaluate the potential of **CB**/**CBT** genes to serve as prognostic biomarkers of **LUAD**, we analyzed clinical data from the **TCGA** to examine if their transcriptional activation in **LUAD** correlated with reduced survival of the patient. The results showed that the expression of 4 **CB** and 2 **CBT** genes in **LUAD** was significantly associated with shorter patient survival (Logrank *p*-value <0.01; **Fig. 9A**).

**Figure 9.**
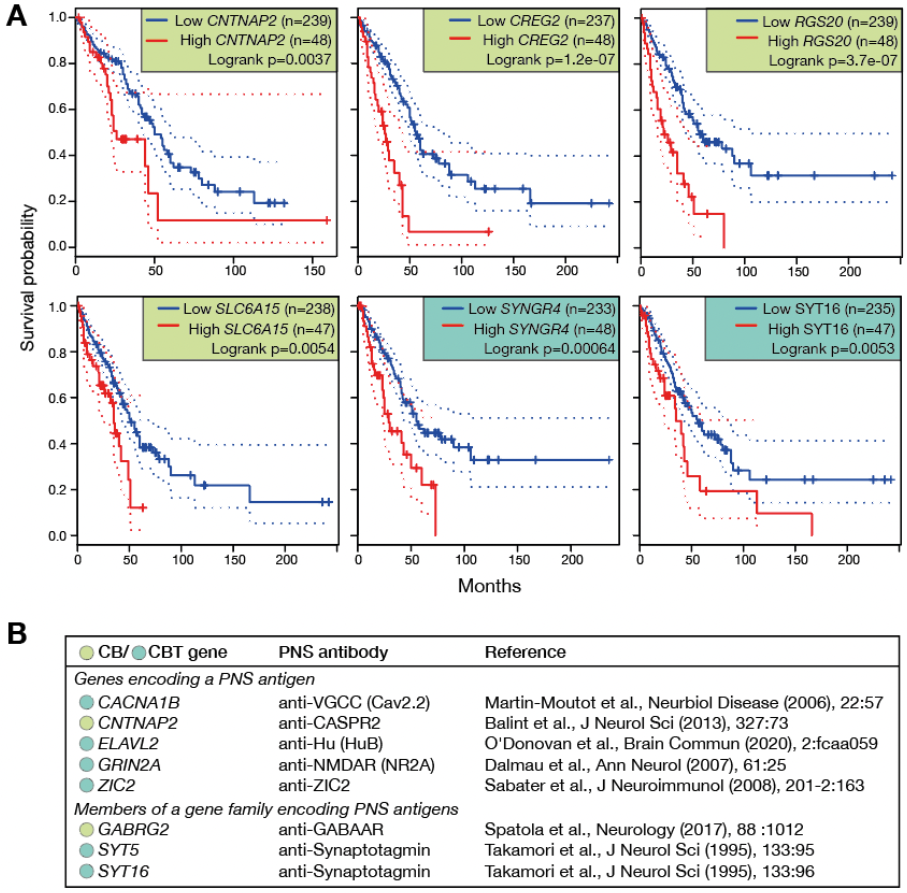
Clinical parameters associated with expression of CB/CBT genes in tumors. **A)** Survival curves (Kaplan-Meier) of **LUAD** patients (**TCGA**) according to high or low m**RNA** levels of the indicated **CB**/**CBT** gene (dotted lines indicate 95% confidence interval). Only **CB**/**CBT** genes with a significant log-rank (p < 0.01) are displayed. **B)** A systematic analysis of the literature was performed to identify **CB**/**CBT** genes that were previously shown to encode PNS-related antigens. Corresponding antibody names and articles references are indicated.

Paraneoplastic neurological syndromes (**PNS**) include a range of cancer-associated autoimmune disorders, which result from the generation of antibodies directed against antigens expressed by both the tumor and the nervous system in the patient. Although **PNS**s occur in less than 0.01% of cancer patients, they are clinically important because neurological manifestations are usually diagnosed well before the underlying tumor. Lung adenocarcinoma belongs to the tumor types that were reported to give rise to **PNS**-inducing autoantibodies (42, 43). About 30 **PNS**-inducing onconeural antigens have been identified to date (44). We decided to examine the contribution of **CB** and **CBT** genes in producing these antigens. Interestingly, five **CB**/**CBT** genes corresponded to genes known to encode **PNS**-associated antigens, and three represented members of gene families reported to encode **PNS**-associated antigens (**Fig. 9B**). This observation demonstrates that activation of **CB**/**CBT** genes in tumors contributes to the production of autoimmune-inducing onconeural proteins.

## Discussion

Cancer development is associated with alterations of transcription networks, which can lead to unscheduled activation of tissue-specific gene expression programs. In the present study, we show that transcriptional changes in **LUAD** include the aberrant expression of a substantial number of brain-expressed genes. As our screening procedure integrated transcriptomic data of **LUAD** tissues and cell lines, as well as **AT2** cells from which **LUAD** originates, expression of these genes in tumoral tissues most likely results from a process of transcriptional activation that occurred in the cancer cells during malignant development. We further provide evidence that many of the brain-expressed genes we identified in **LUAD** also become activated in other types of tumors.

Ectopic activation of brain-expressed genes in tumors has been reported previously, but most often on a gene by gene basis (45-47). Our present study shows that brain-expressed genes actually represent a substantial proportion of the genes that become aberrantly activated in **LUAD** tumors, suggesting engagement of a brain transcriptional circuit in these tumors. Our transcriptomic analyses did not identify subgroups of **LUAD** tumors showing convergent expression of multiple **CB**/**CBT** genes, but rather revealed scattered activation of these genes in most tumor samples. This suggests that **LUAD** tumors share a mechanism of transcriptional dysregulation that does not necessarily impose, but only facilitates activation of **CB**/**CBT** genes. Our observations suggest that impaired activity of the **REST** transcriptional repressor represents a major route to the weakening of **CB**/**CBT** gene silencing in tumor cells. The **REST** factor is known to play a key role in the repression of neuronal genes in non-neuronal cells (48). Consistently, most **CB** and **CBT** genes were expressed in neuronal cells in the brain. Interestingly, previous studies have reported impaired **REST** activities in tumors of non-neuronal origin (48, 49). Multiple mechanisms appear contribute to reduce **REST** functions in cancer cells, including transcriptional downregulation, mutation, alternative splicing, or loss of function of co-factors (50). This may explain our observation that activation of **CB**/**CBT** genes was often, albeit not always, associated with reduced ***REST*** m**RNA** levels in tumor cell lines. In addition, it is possible the that tumoral activation of several **CB**/**CBT** genes results from mechanisms that are independent of **REST**.

Impaired **REST** activities have been shown to promote the development of tumors of non-neural origin, but the underlying mechanisms remain partially understood (48, 50). It is reasonable to propose that these include de-repression of neuronal genes with oncogenic functions, and that several **CB**/**CBT** genes may be part of these **REST**-regulated pro-tumorigenic mediators. A quick survey of the CancerMine literature-mined database identified several **CB** and **CBT** genes with potential oncogenic functions. ***POU3F2***, for instance, was found to be upregulated in malignant melanoma where it represses the melanocyte lineage determining transcription factor MITF, thereby favoring invasiveness (51). ***KCNK9***, which encodes a potassium channel regulating neuronal cell excitability, was shown to promote resistance to hypoxia and serum deprivation in tumors where it is upregulated (52). ***GPR19***, a gene encoding an orphan G-protein-coupled receptor involved in circadian rhythm regulation in the brain, was shown to promote cell proliferation by accelerating the **G2/M** transition of the cell cycle in lung tumor cells that express the gene (53, 54). As a last example, ***ZIC2*** encodes a transcriptional regulator involved in brain organogenesis, and its transcriptional activation in liver cancer was found to drive the self-renewal of cancer stem cells (55).

Beyond the fact that individual brain-expressed genes can have oncogenic potential, the observation that a substantial amount of **CB**/**CBT** genes become expressed in **LUAD** raises the intriguing possibility that activation of an integrated network of neuronal functions in cancer cells might contribute to malignant progression. In line with this assumption, is the increasing evidence that supports existence of a communication framework between cancers and nerves (56, 57). Several tumors for instance, were shown to use surrounding nerves as a route of dissemination, through a process of “perineural invasion”. Other studies suggest that axons innervating the tumors release molecules that modulate signaling processes in cancer cells, which, for instance, induce cell proliferation. Conversely, cancer cells were reported to secrete molecules that stimulate the outgrowth of nerves into the tumor, a process termed neo-neurogenesis. Our hypothesis is that **CB**/**CBT** genes expressed in the cancer cells contribute to establishing molecular networks that allows communications with neuronal cells.

Paraneoplastic neurological syndromes (**PNS**) are rare autoimmune disorders occurring in association with active or subclinical early-stage cancers. Manifestation of **PNS** often precedes diagnosis of the tumor, and can therefore help to set up an early follow-up of the cancer patient (58). The discovery of anti-neuronal autoantibodies in **PNS** patients has allowed a major advance in the understanding of these syndromes (44). The mechanisms underlying expression of neuronal antigens by the tumor cells remain however unclear. We found that a remarkable proportion of **CB**/**CBT** genes encode previously described **PNS**-related antigens. Since many **PNS**-inducing antigens remain unknown (58), exploration of the **CB**/**CBT** genes we identified in the present study may help to characterize new such antigens. This may be useful for the development of early diagnosis of **PNS**. Interestingly, most of the **CB**/**CBT** genes that encode known **PNS**-related antigens (4/5) were predicted to be regulated by **REST**. This suggests that **PNS** may in some cases be a manifestation of the presence of a tumor with reduced **REST** activities.

## Acknowledgments

This work was supported by grants from the Fonds de la recherche scientifique (FRS-FNRS), Belgium (ref. J.0032.19), the D.G. Higher Education and Scientific Research of the French Community of Belgium (Action de Recherches Concertées), and the Fonds special de recherche (FSR) of the UCLouvain, Belgium. A.D. was supported by a fellowship of the FRS-FNRS-FRIA, Belgium (ref. 1.E008.19), and by a grant of the health science sector of UCLouvain (Bourse du patrimoine, 2021). A.L. was supported by the de Duve Institute, Brussels, Belgium.

## Notes

### Competing Interest Statement

The authors have declared no competing interest.

## References

1 Koslowski M, Bell C, Seitz G, Lehr HA, Roemer K, Muntefering H, Huber C, Sahin U and Tureci O: Frequent nonrandom activation of germ-line genes in human cancer. Cancer Res 64(17): 5988–5993., 2004. PMID, DOI:

2 De Smet C and Loriot A: DNA hypomethylation and activation of germline-specific genes in cancer. Adv Exp Med Biol 754(149–166, 2013. PMID, DOI:

3 Axelsen JB, Lotem J, Sachs L and Domany E: Genes overexpressed in different human solid cancers exhibit different tissue-specific expression profiles. Proc Natl Acad Sci U S A 104(32): 13122–13127, 2007. PMID: PMC1941809, DOI: 10.1073/pnas.0705824104

4 Diacofotaki A, Loriot A and De Smet C: Identification of tissue-specific gene clusters induced by DNA demethylation in lung adenocarcinoma: More than germline genes. Cancers (Basel) 14(4): 1007, 2022. PMID, DOI: 10.3390/cancers14041007

5 Lotem J, Netanely D, Domany E and Sachs L: Human cancers overexpress genes that are specific to a variety of normal human tissues. Proc Natl Acad Sci U S A 102(51): 18556–18561, 2005. PMID: PMC1317977, DOI: 10.1073/pnas.0509360102

6 Rousseaux S, Debernardi A, Jacquiau B, Vitte AL, Vesin A, Nagy-Mignotte H, Moro-Sibilot D, Brichon PY, Lantuejoul S, Hainaut P, Laffaire J, de Reynies A, Beer DG, Timsit JF, Brambilla C, Brambilla E and Khochbin S: Ectopic activation of germline and placental genes identifies aggressive metastasis-prone lung cancers. Sci Transl Med 5(186): 186ra166, 2013. PMID: PMC4818008, DOI: 10.1126/scitranslmed.3005723

7 Shukla S, Cyrta J, Murphy DA, Walczak EG, Ran L, Agrawal P, Xie Y, Chen Y, Wang S, Zhan Y, Li D, Wong EWP, Sboner A, Beltran H, Mosquera JM, Sher J, Cao Z, Wongvipat J, Koche RP, Gopalan A, Zheng D, Rubin MA, Scher HI, Chi P and Chen Y: Aberrant activation of a gastrointestinal transcriptional circuit in prostate cancer mediates castration resistance. Cancer Cell 32(6): 792–806 e797, 2017. PMID: PMC5728174, DOI: 10.1016/j.ccell.2017.10.008

8 Akers SN, Odunsi K and Karpf AR: Regulation of cancer germline antigen gene expression: Implications for cancer immunotherapy. Future Oncol 6(5): 717–732, 2010. PMID, DOI:

9 Visvader JE: Cells of origin in cancer. Nature 469(7330): 314–322, 2011. PMID, DOI: 10.1038/nature09781

10 Blanpain C: Tracing the cellular origin of cancer. Nat Cell Biol 15(2): 126–134, 2013. PMID, DOI: 10.1038/ncb2657

11 Wang Z, Li Z, Zhou K, Wang C, Jiang L, Zhang L, Yang Y, Luo W, Qiao W, Wang G, Ni Y, Dai S, Guo T, Ji G, Xu M, Liu Y, Su Z, Che G and Li W: Deciphering cell lineage specification of human lung adenocarcinoma with single-cell rna sequencing. Nat Commun 12(1): 6500, 2021. PMID: PMC8586023, DOI: 10.1038/s41467-021-26770-2

12 Whitsett JA, Kalin TV, Xu Y and Kalinichenko VV: Building and regenerating the lung cell by cell. Physiol Rev 99(1): 513–554, 2019. PMID: PMC6442926, DOI: 10.1152/physrev.00001.2018

13 Weeden CE, Chen Y, Ma SB, Hu Y, Ramm G, Sutherland KD, Smyth GK and Asselin-Labat ML: Lung basal stem cells rapidly repair DNA damage using the error-prone nonhomologous end-joining pathway. PLoS Biol 15(1): e2000731, 2017. PMID: PMC5268430, DOI: 10.1371/journal.pbio.2000731

14 Zuber V, Marconett CN, Shi J, Hua X, Wheeler W, Yang C, Song L, Dale AM, Laplana M, Risch A, Witoelar A, Thompson WK, Schork AJ, Bettella F, Wang Y, Djurovic S, Zhou B, Borok Z, van der Heijden HF, de Graaf J, Swinkels D, Aben KK, McKay J, Hung RJ, Bikeboller H, Stevens VL, Albanes D, Caporaso NE, Han Y, Wei Y, Panadero MA, Mayordomo JI, Christiani DC, Kiemeney L, Andreassen OA, Houlston R, Amos CI, Chatterjee N, Laird-Offringa IA, Mills IG and Landi MT: Pleiotropic analysis of lung cancer and blood triglycerides. J Natl Cancer Inst 108(12), 2016. PMID: PMC5241892, DOI: 10.1093/jnci/djw167

15 Bolger AM, Lohse M and Usadel B: Trimmomatic: A flexible trimmer for illumina sequence data. Bioinformatics 30(15): 2114–2120, 2014. PMID: PMC4103590, DOI: 10.1093/bioinformatics/btu170

16 Kim D, Paggi JM, Park C, Bennett C and Salzberg SL: Graph-based genome alignment and genotyping with hisat2 and hisat-genotype. Nat Biotechnol 37(8): 907–915, 2019. PMID: PMC7605509, DOI: 10.1038/s41587-019-0201-4

17 Liao Y, Smyth GK and Shi W: Featurecounts: An efficient general purpose program for assigning sequence reads to genomic features. Bioinformatics 30(7): 923–930, 2014. PMID, DOI: 10.1093/bioinformatics/btt656

18 Krueger F and Andrews SR: Bismark: A flexible aligner and methylation caller for bisulfite-seq applications. Bioinformatics 27(11): 1571–1572, 2011. PMID: PMC3102221, DOI: 10.1093/bioinformatics/btr167

19 Consortium GT: The genotype-tissue expression (gtex) project. Nat Genet 45(6): 580–585, 2013. PMID: PMC4010069, DOI: 10.1038/ng.2653

20 Diacofotaki A, Loriot A and De Smet C: Identification of tissue-specific gene clusters induced by DNA demethylation in lung adenocarcinoma: More than germline genes. Cancers (Basel) 14(4), 2022. PMID: PMC8870412, DOI: 10.3390/cancers14041007

21 Suzuki A, Makinoshima H, Wakaguri H, Esumi H, Sugano S, Kohno T, Tsuchihara K and Suzuki Y: Aberrant transcriptional regulations in cancers: Genome, transcriptome and epigenome analysis of lung adenocarcinoma cell lines. Nucleic Acids Res 42(22): 13557–13572, 2014. PMID: PMC4267666, DOI: 10.1093/nar/gku885

22 Fain JS, Loriot A, Diacofotaki A, Van Tongelen A and De Smet C: Transcriptional overlap links DNA hypomethylation with DNA hypermethylation at adjacent promoters in cancer. Sci Rep 11(1): 17346, 2021. PMID: PMC8405634, DOI: 10.1038/s41598-021-96844-0

23 Cancer Genome Atlas Research N: Comprehensive molecular profiling of lung adenocarcinoma. Nature 511(7511): 543–550, 2014. PMID: PMC4231481, DOI: 10.1038/nature13385

24 Colaprico A, Silva TC, Olsen C, Garofano L, Cava C, Garolini D, Sabedot TS, Malta TM, Pagnotta SM, Castiglioni I, Ceccarelli M, Bontempi G and Noushmehr H: Tcgabiolinks: An r/bioconductor package for integrative analysis of tcga data. Nucleic Acids Res 44(8): e71, 2016. PMID: PMC4856967, DOI: 10.1093/nar/gkv1507

25 Yu G, Wang LG, Han Y and He QY: Clusterprofiler: An r package for comparing biological themes among gene clusters. OMICS 16(5): 284–287, 2012. PMID: PMC3339379, DOI: 10.1089/omi.2011.0118

26 Robinson JT, Thorvaldsdottir H, Winckler W, Guttman M, Lander ES, Getz G and Mesirov JP: Integrative genomics viewer. Nat Biotechnol 29(1): 24–26, 2011. PMID: PMC3346182, DOI: 10.1038/nbt.1754

27 Uhlen M, Bjorling E, Agaton C, Szigyarto CA, Amini B, Andersen E, Andersson AC, Angelidou P, Asplund A, Asplund C, Berglund L, Bergstrom K, Brumer H, Cerjan D, Ekstrom M, Elobeid A, Eriksson C, Fagerberg L, Falk R, Fall J, Forsberg M, Bjorklund MG, Gumbel K, Halimi A, Hallin I, Hamsten C, Hansson M, Hedhammar M, Hercules G, Kampf C, Larsson K, Lindskog M, Lodewyckx W, Lund J, Lundeberg J, Magnusson K, Malm E, Nilsson P, Odling J, Oksvold P, Olsson I, Oster E, Ottosson J, Paavilainen L, Persson A, Rimini R, Rockberg J, Runeson M, Sivertsson A, Skollermo A, Steen J, Stenvall M, Sterky F, Stromberg S, Sundberg M, Tegel H, Tourle S, Wahlund E, Walden A, Wan J, Wernerus H, Westberg J, Wester K, Wrethagen U, Xu LL, Hober S and Ponten F: A human protein atlas for normal and cancer tissues based on antibody proteomics. Mol Cell Proteomics 4(12): 1920–1932, 2005. PMID, DOI: 10.1074/mcp.M500279-MCP200

28 Drouin-Ouellet J, Lau S, Brattas PL, Rylander Ottosson D, Pircs K, Grassi DA, Collins LM, Vuono R, Andersson Sjoland A, Westergren-Thorsson G, Graff C, Minthon L, Toresson H, Barker RA, Jakobsson J and Parmar M: Rest suppression mediates neural conversion of adult human fibroblasts via microrna-dependent and -independent pathways. EMBO Mol Med 9(8): 1117–1131, 2017. PMID: PMC5538296, DOI: 10.15252/emmm.201607471

29 Warren A, Chen Y, Jones A, Shibue T, Hahn WC, Boehm JS, Vazquez F, Tsherniak A and McFarland JM: Global computational alignment of tumor and cell line transcriptional profiles.Nat Commun 12(1): 22, 2021. PMID: PMC7782593, DOI: 10.1038/s41467-020-20294-x

30 Tang Z, Kang B, Li C, Chen T and Zhang Z: Gepia2: An enhanced web server for large-scale expression profiling and interactive analysis. Nucleic Acids Res 47(W1): W556–W560, 2019. PMID: PMC6602440, DOI: 10.1093/nar/gkz430

31 Gu Z, Eils R and Schlesner M: Complex heatmaps reveal patterns and correlations in multidimensional genomic data. Bioinformatics 32(18): 2847–2849, 2016. PMID, DOI: 10.1093/bioinformatics/btw313

32 Loriot A, Van Tongelen A, Blanco J, Klaessens S, Cannuyer J, van Baren N, Decottignies A and De Smet C: A novel cancer-germline transcript carrying pro-metastatic mir-105 and tet-targeting mir-767 induced by DNA hypomethylation in tumors. Epigenetics 9(8): 1163–1171, 2014. PMID: PMC4164501, DOI: 10.4161/epi.29628

33 Uhlen M, Fagerberg L, Hallstrom BM, Lindskog C, Oksvold P, Mardinoglu A, Sivertsson A, Kampf C, Sjostedt E, Asplund A, Olsson I, Edlund K, Lundberg E, Navani S, Szigyarto CA, Odeberg J, Djureinovic D, Takanen JO, Hober S, Alm T, Edqvist PH, Berling H, Tegel H, Mulder J, Rockberg J, Nilsson P, Schwenk JM, Hamsten M, von Feilitzen K, Forsberg M, Persson L, Johansson F, Zwahlen M, von Heijne G, Nielsen J and Ponten F: Proteomics. Tissue-based map of the human proteome. Science 347(6220): 1260419, 2015. PMID, DOI: 10.1126/science.1260419

34 Bhattacharjee A, Richards WG, Staunton J, Li C, Monti S, Vasa P, Ladd C, Beheshti J, Bueno R, Gillette M, Loda M, Weber G, Mark EJ, Lander ES, Wong W, Johnson BE, Golub TR, Sugarbaker DJ and Meyerson M: Classification of human lung carcinomas by mrna expression profiling reveals distinct adenocarcinoma subclasses. Proc Natl Acad Sci U S A 98(24): 13790–13795, 2001. PMID: PMC61120, DOI: 10.1073/pnas.191502998

35 Kosari F, Ida CM, Aubry MC, Yang L, Kovtun IV, Klein JL, Li Y, Erdogan S, Tomaszek SC, Murphy SJ, Bolette LC, Kolbert CP, Yang P, Wigle DA and Vasmatzis G: Ascl1 and ret expression defines a clinically relevant subgroup of lung adenocarcinoma characterized by neuroendocrine differentiation. Oncogene 33(29): 3776–3783, 2014. PMID: PMC4329973, DOI: 10.1038/onc.2013.359

36 Borges M, Linnoila RI, van de Velde HJ, Chen H, Nelkin BD, Mabry M, Baylin SB and Ball DW: An achaete-scute homologue essential for neuroendocrine differentiation in the lung. Nature 386(6627): 852–855, 1997. PMID, DOI: 10.1038/386852a0

37 Borromeo MD, Savage TK, Kollipara RK, He M, Augustyn A, Osborne JK, Girard L, Minna JD, Gazdar AF, Cobb MH and Johnson JE: Ascl1 and neurod1 reveal heterogeneity in pulmonary neuroendocrine tumors and regulate distinct genetic programs. Cell Rep 16(5): 1259–1272, 2016. PMID: PMC4972690, DOI: 10.1016/j.celrep.2016.06.081

38 Casarosa S, Fode C and Guillemot F: Mash1 regulates neurogenesis in the ventral telencephalon. Development 126(3): 525–534, 1999. PMID, DOI: 10.1242/dev.126.3.525

39 Schoenherr CJ and Anderson DJ: The neuron-restrictive silencer factor (nrsf): A coordinate repressor of multiple neuron-specific genes. Science 267(5202): 1360–1363, 1995. PMID, DOI: 10.1126/science.7871435

40 Cortes-Sarabia K, Alarcon-Romero LDC, Flores-Alfaro E, Illades-Aguiar B, Vences-Velazquez A, Mendoza-Catalan MA, Navarro-Tito N, Valdes J, Moreno-Godinez ME and Ortuno-Pineda C: Significant decrease of a master regulator of genes (rest/nrsf) in high-grade squamous intraepithelial lesion and cervical cancer. Biomed J 44(6 Suppl 2): S171–S178, 2021. PMID: PMC9068566, DOI: 10.1016/j.bj.2020.08.012

41 Gurrola-Diaz C, Lacroix J, Dihlmann S, Becker CM and von Knebel Doeberitz M: Reduced expression of the neuron restrictive silencer factor permits transcription of glycine receptor alpha1 subunit in small-cell lung cancer cells. Oncogene 22(36): 5636–5645, 2003. PMID, DOI: 10.1038/sj.onc.1206790

42 Banjara M, Ghosh C, Dadas A, Mazzone P and Janigro D: Detection of brain-directed autoantibodies in the serum of non-small cell lung cancer patients. PLoS One 12(7): e0181409, 2017. PMID: PMC5528996, DOI: 10.1371/journal.pone.0181409

43 Ma J, Wang A, Jiang W, Ma L and Lin Y: Clinical characteristics of paraneoplastic neurological syndrome related to different pathological lung cancers. Thorac Cancer 12(16): 2265–2270, 2021. PMID: PMC8364989, DOI: 10.1111/1759-7714.14070

44 Lancaster E and Dalmau J: Neuronal autoantigens--pathogenesis, associated disorders and antibody testing. Nat Rev Neurol 8(7): 380–390, 2012. PMID: PMC3718498, DOI: 10.1038/nrneurol.2012.99

45 Grelet S, Andries V, Polette M, Gilles C, Staes K, Martin AP, Kileztky C, Terryn C, Dalstein V, Cheng CW, Shen CY, Birembaut P, Van Roy F and Nawrocki-Raby B: The human nanos3 gene contributes to lung tumour invasion by inducing epithelial-mesenchymal transition. J Pathol 237(1): 25–37, 2015. PMID, DOI: 10.1002/path.4549

46 Martin NL, Saba-El-Leil Mk, Sadekova S, Meloche S and Sauvageau G: En2 is a candidate oncogene in human breast cancer. Oncogene 24(46): 6890–6901, 2005. PMID, DOI: 10.1038/sj.onc.1208840

47 Zong X, Yang H, Yu Y, Zou D, Ling Z, He X and Meng X: Possible role of pax-6 in promoting breast cancer cell proliferation and tumorigenesis. BMB Rep 44(9): 595–600, 2011. PMID, DOI: 10.5483/bmbrep.2011.44.9.595

48 Coulson JM: Transcriptional regulation: Cancer, neurons and the rest. Curr Biol 15(17): R665–668, 2005. PMID, DOI: 10.1016/j.cub.2005.08.032

49 Wagoner MP, Gunsalus KT, Schoenike B, Richardson AL, Friedl A and Roopra A: The transcription factor rest is lost in aggressive breast cancer. PLoS Genet 6(6): e1000979, 2010. PMID: PMC2883591, DOI: 10.1371/journal.pgen.1000979

50 Huang Z and Bao S: Ubiquitination and deubiquitination of rest and its roles in cancers. FEBS Lett 586(11): 1602–1605, 2012. PMID: PMC3361610, DOI: 10.1016/j.febslet.2012.04.052

51 Goodall J, Carreira S, Denat L, Kobi D, Davidson I, Nuciforo P, Sturm RA, Larue L and Goding CR: Brn-2 represses microphthalmia-associated transcription factor expression and marks a distinct subpopulation of microphthalmia-associated transcription factor-negative melanoma cells. Cancer Res 68(19): 7788–7794, 2008. PMID, DOI: 10.1158/0008-5472.CAN-08-1053

52 Mu Y and Gage FH: Adult hippocampal neurogenesis and its role in alzheimer’s disease. Mol Neurodegener 6(85, 2011. PMID: 3261815, DOI: 10.1186/1750-1326-6-85

53 Kastner S, Voss T, Keuerleber S, Glockel C, Freissmuth M and Sommergruber W: Expression of g protein-coupled receptor 19 in human lung cancer cells is triggered by entry into s-phase and supports g(2)-m cell-cycle progression. Mol Cancer Res 10(10): 1343–1358, 2012. PMID, DOI: 10.1158/1541-7786.MCR-12-0139

54 Yamaguchi Y, Murai I, Goto K, Doi S, Zhou H, Setsu G, Shimatani H, Okamura H, Miyake T and Doi M: Gpr19 is a circadian clock-controlled orphan gpcr with a role in modulating free-running period and light resetting capacity of the circadian clock. Sci Rep 11(1): 22406, 2021. PMID: PMC8599615, DOI: 10.1038/s41598-021-01764-8

55 Zhu P, Wang Y, He L, Huang G, Du Y, Zhang G, Yan X, Xia P, Ye B, Wang S, Hao L, Wu J and Fan Z: Zic2-dependent oct4 activation drives self-renewal of human liver cancer stem cells. J Clin Invest 125(10): 3795–3808, 2015. PMID: PMC4607118, DOI: 10.1172/JCI81979

56 Jobling P, Pundavela J, Oliveira SM, Roselli S, Walker MM and Hondermarck H: Nerve-cancer cell cross-talk: A novel promoter of tumor progression. Cancer Res 75(9): 1777–1781, 2015. PMID, DOI: 10.1158/0008-5472.CAN-14-3180

57 Wang H, Zheng Q, Lu Z, Wang L, Ding L, Xia L, Zhang H, Wang M, Chen Y and Li G: Role of the nervous system in cancers: A review. Cell Death Discov 7(1): 76, 2021. PMID: PMC8041826, DOI: 10.1038/s41420-021-00450-y

58 Honnorat J and Antoine JC: Paraneoplastic neurological syndromes. Orphanet J Rare Dis 2(22, 2007. PMID: PMC1868710, DOI: 10.1186/1750-1172-2-22

